# HER2 mutation–derived neoantigens in NSCLC as actionable targets for TCR therapy

**DOI:** 10.64898/2026.07.30.741830

**Authors:** Amanda Montoya, Hui Nie, Peixin Jiang, Jared Slone, Yulia Shulga, Anika Patel, Prashant Menon, Isabella Polic, Emily Bontekoe, Lingzhi Hong, Minying Zhang, Emane Rose Assita, Sarah Forward, Changsheng Xing, Bo Jiang, Drew C. Deniger, Gregory A. Lizee, Maura Gillison, Navin Varadarajan, Xiuning Le, Jianjun Zhang, Lydia Kavraki, Sheldon J.J. Kwok, John V. Heymach, Alexandre Reuben

## Abstract

HER2 mutations are oncogenic drivers in 1–6% of non-small cell lung cancers (NSCLC), but therapeutic resistance limits the durability of current HER2-targeted treatments. Here, we identify T-cell receptors (TCRs) targeting recurrent HER2 hotspot mutations as a potential immunotherapeutic strategy for HER2-mutant NSCLC. Using neoepitope prediction and antigen-specific T-cell enrichment, we isolated HLA-A*02:01restricted TCRs recognizing HER2 A775insYVMA, S310F, and G776delinsVC mutations, collectively covering approximately 60% of HER2-mutant NSCLC. These TCRs selectively recognized mutant HER2 epitopes without detectable wild-type reactivity and some displayed cross-recognition of related hotspot variants, expanding the spectrum of targetable tumors. The G776delinsVC-specific TCR also exhibited co-receptor-independent activity showcased by its ability to activate CD4^+^ T cells. Importantly, timelapse single-cell flow cytometry analyses demonstrated that TCR-engineered T cells repeatedly reacquired activated polyfunctional states following serial antigen stimulation, while serial tumor rechallenge assays confirmed sustained cytotoxic activity across multiple rounds of tumor killing. These findings identify recurrent HER2 mutations as shared immunotherapeutic targets and provide a foundation for the development of TCR-based therapies for HER2-mutant NSCLC.

## Introduction

The human epidermal growth factor receptor 2 (HER2, ERBB2) is a receptor tyrosine kinase that regulates cell proliferation, survival, and differentiation (1). Aberrant HER2 signaling is a well-established oncogenic driver across multiple tumor types, mediated through gene amplification, protein overexpression, or activating mutations (2). In non– small cell lung cancer (NSCLC), HER2 alterations occur in approximately 1–6% of cases and are prevalent among women without prior smoking history. Exon 20 insertions, mutations occurring within the tyrosine kinase domain, are the most prevalent mutations in NSCLC; although point mutations and less frequent structural variants have also been described, contributing to biologic and therapeutic heterogeneity in this setting. These tumors are frequently associated with aggressive clinical behavior, including a high incidence of brain metastases (3, 4).

A range of HER2-targeted strategies including monoclonal antibodies, tyrosine kinase inhibitors, and antibody–drug conjugates have broadened treatment options for patients with HER2-mutant NSCLC. Trastuzumab deruxtecan, a HER2-targeted antibody–drug conjugate, has shown consistent clinical activity in HER2-mutant tumors and remains a central therapeutic option in this setting (5). More recently, the treatment landscape for HER2-positive NSCLC broadened with the approval of zongertinib and sevabertinib, two selective HER2 kinase inhibitors for previously treated tumors with HER2 tyrosine kinase domain mutations (6, 7). Nevertheless, the clinical impact of these therapies has been constrained by the eventual development of resistance, which reinforces the need for more mutation-directed therapeutics (8–10). Importantly, most currently approved agents target the kinase domain or extracellular receptor irrespective of the specific activating mutation present, and resistance mechanisms frequently involve secondary alterations or adaptive signaling pathways (9, 10).

A promising therapeutic strategy for HER2-positive NSCLC is adoptive T-cell therapy (ACT). ACT encompasses a range of approaches in which patient-derived T cells are expanded or engineered *ex vivo* to improve their capacity to target malignant tumor cells (11). Tumor-infiltrating lymphocyte (TIL) therapy has demonstrated promising responses in NSCLC. However, its clinical benefit remains inconsistent, likely because of heterogeneity in TIL populations across patients (12, 13). Chimeric antigen receptor (CAR) T cells represent an engineered T-cell approach in which tumor recognition is redirected toward specific cell-surface antigens through the introduction of a synthetic receptor capable of recognizing these antigens (14). Although HER2-targeted CAR T cells have demonstrated potent antitumor activity in preclinical models, early clinical translation was hindered by severe on-target/off-tumor associated toxicities after infusion of high-affinity HER2 targeting CAR T cells (15–17). These events underscore the difficulty of safely targeting this widely expressed surface receptor. In contrast, T cell receptor-engineered T cells (TCR-T) can recognize intracellularly derived peptides presented on human leukocyte antigen (HLA) molecules, permitting selective targeting of tumorspecific antigens that are not accessible to CAR-based approaches (18).

Somatic mutations can give rise to neoantigens, which are processed and presented as mutated peptides that are, by definition, not found in normal tissues (19, 20). Hotspot mutations represent attractive targets because they generate recurrent, tumor-restricted epitopes that can be recognized across many patients, supporting the development of broadly applicable T-cell therapies (21–23). Several recurrent HER2 hotspot mutations, including exon 20 insertions, are shared across patients, supporting the feasibility of developing comprehensive, HLA-restricted TCR therapies (3, 4). In addition to point mutations, insertion–deletion events (indels) can also give rise to immunogenic neoantigens by creating novel peptide sequences not present in the human proteome (24). This is particularly important in HER2-driven NSCLC, where recurrent HER2 mutations frequently involve indels, such as the YVMA insertion at 772 and the G776delinsVC variant. By redirecting T cells against mutated HER2 peptides rather than the broadly expressed wild-type receptor, mutation-specific HER2 TCR-T therapy offers the potential for potent tumor recognition with a substantially improved safety profile. Notably, mutation-specific TCR therapies targeting oncogenic drivers such as KRAS in NSCLC have demonstrated objective tumor regressions, including partial responses, supporting the clinical feasibility of this approach (25).

Although HER2 mutations are increasingly recognized as actionable drivers in NSCLC, there remains a significant gap in the development of T-cell receptor–based therapies capable of targeting these alterations. To date, the only reported HER2-directed TCRs have been restricted to class II epitopes (26), and no class I–restricted, mutation-specific HER2 TCRs have been functionally validated in the context of NSCLC. Here, we present three HLA-A*02:01–restricted class I TCRs directed against common hotspot HER2 mutations found in NSCLC, which together account for ∼60% of all HER2 hotspot mutations in this disease. We functionally validate their specificity, cytotoxic activity, and absence of detectable reactivity against the normal human proteome while demonstrating HER2 mutation degeneracy for S310F/Y and G776delinsVC/IC/IV. This work establishes a foundational framework for mutation-specific HER2 TCR-T therapies in NSCLC and broadens the therapeutic landscape beyond previously reported class II– restricted targets.

## Methods

### ERBB2 Mutation Analysis

Somatic mutation data were obtained from the TCGA PanCancer Atlas cohort through cBioPortal (27). ERBB2 mutations were extracted from downloaded mutation annotation files and analyzed using custom scripts written in Python 3. Amino acid positions were derived from protein-level mutation annotations, and recurrent alterations were quantified by frequency within the analyzed cohort. Lollipop plots depicting the distribution of ERBB2 mutations across the protein sequence were generated using the Python libraries pandas and matplotlib. Clonality classifications were obtained from the published Guardant360 cohort reported by Hong et al (3). In the original study, variant clonality was determined by normalizing variant allele frequency (VAF) to the maximum somatic VAF detected within each sample, with variants classified as clonal when the normalized VAF was ≥0.5. Previously reported clonality assignments were used for data visualization in the present study.

### Cell lines

H1975 cells were purchased from the American Type Culture Collection (ATCC). H1781, 293GP, and Lenti293 cell lines were obtained from the laboratories of Dr. John Heymach and Dr. Maura Gillison at the MD Anderson Cancer Center and were authen-ticated by the MD Anderson Cytogenetic and Cell Authentication Core by short tandem repeat (STR) profiling before use.

### Generation of HLA-A*02:01–Expressing Cell Lines

The H1975-A*02:01 cell line was generated by cloning a synthesized HLA-A*02:01 al-lele (GeneScript USA, Inc., Piscataway, NJ) into a pENTER vector, followed by transfer of the final allele construct into the lentiviral vector pLV401GFP using Gateway™ LR Clonase™ II Enzyme Mix (Invitrogen™, 11791020). The H1781-A*02:01 cell line was generated by transduction with a lentiviral vector produced by VectorBuilder (Vector ID: VB250619-2298uvq). Lentiviral supernatants were generated by transfecting Lenti293 packaging cells with the transfer vector and packaging plasmids using Lipofectamine™ 3000 (Thermofisher, L00015) according to the manufacturer’s protocol. Viral supernatants were collected at 48–72 hours and used immediately for transduction. Tumor cell lines successfully transduced with HLA constructs were isolated by fluorescence-activated cell sorting (FACS) based on GFP expression.

### Neoepitope prediction

Potential HER2 mutation–derived neoepitopes were predicted using NetMHCpan-4.1 (28). Candidate peptides were selected based on their predicted high-affinity binding (percent rank < 0.5) to the ten most prevalent HLA class I alleles in the United States (21). Synthetic peptides were produced by GenScript USA, Inc. (Piscataway, NJ) at >95% purity.

### Peripheral blood mononuclear cell (PMBC) isolation

Apheresis products from HLA-A*02:01 positive healthy donors were sourced through Charles River and processed immediately upon receipt. PBMCs were isolated using ficoll density centrifugation and cryopreserved in freezing medium consisting of 90% FBS and 10% DMSO for further use.

### Generation of Monocyte-Derived Dendritic Cells (DCs)

Monocytes were enriched by plastic adherence as previously described (29). Briefly, 1×10 mononuclear cells were plated in T75 flasks with 15 mL CTS™ AIM V™ medium (Gibco, A3830801) and incubated at 37°C for 2 h to allow monocyte adherence. Nonadherent cells were gently removed, and flasks were washed twice with PBS. Adherent monocytes were cultured for 72 h in CTS™ AIM V™ medium supplemented with 800 IU/mL GM-CSF (Peprotech, 300-03) and 400 IU/mL IL-4 (PeproTech, 200-04). To induce maturation, cells were stimulated overnight with 10 ng/mL LPS and 100 IU/mL IL-2 (R&D Systems, BT-002-AFL-050). Mature DCs were harvested by vigorous resuspension and pulsed with 10 µg/mL peptide at 37°C for 2 h. Prior to co-culture with T cells, DCs were irradiated with 30 Gy to prevent expansion of contaminating NK cells or memory T cells.

### Preparation of naïve CD8^⁺^ T cells

CD8⁺ T cells were isolated from HLA-A*02:01 positive PBMCs using the EasySep™ Human CD8⁺ Negative Isolation Kit (STEMCELL Technologies, 17953). Purified CD8⁺ T cells were rested overnight in LymphoONE™ T-Cell Expansion Xeno-Free Medium (Takara, WK552) supplemented with 5% human serum and 5 ng/mL IL-7 (Peprotech, 200-07).

### Antigen specific T cell generation

DCs and naïve CD8⁺ T cells were counted and resuspended in LymphoONE™ T-Cell Expansion Xeno-Free Medium containing 5% human serum at 5×10 cells/mL and 2×10 cells/mL, respectively. IL-21 (60 ng/mL; PeproTech 200-21) was added to the Tcell suspension. DCs and T cells were combined at a 1:1 (vol/vol) ratio, resulting in an effective 4:1 T cell:DC ratio. A total of 500 µL of the mixture was added to each well of a 48-well plate and incubated at 37°C for 72 h. IL-15 (Peprotech, 200-15) and IL-7 (5 ng/mL each; final concentration) were then added to each well, and cultures were incubated for an additional 72 h. Cells were transferred to 12-well plates with fresh medium, IL-7, and IL-15, and maintained until analysis on day 10.

### Flow Cytometry and Cell Sorting

Anti-human CD8-APC (clone SK1), CD4-BB700 (OKT4), mouse TCRβ-FITC (H57-597) were purchased from BioLegend (344722, 317416 and 19206 respectively). PE-labeled peptide–Major Histocompatibility Complex (MHC) tetramers were produced by the MHC Tetramer Production Facility (Baylor College of Medicine). For sorting, cells were harvested, washed with FACS buffer, and resuspended at 1–50 × 10 cells/mL. Fluorophore-conjugated antibodies were added and incubated for 30 min at 4°C. After washing, samples were analyzed on Cytek Aurora and sorted on a FACSAria I Cell Sorter (BD Biosciences). Data was analyzed using FlowJo 10.10 software (TreeStar). Cells were sorted for live/CD8⁺/tetramer⁺ populations.

### Identification of αβ TCR Chains

Antigen-reactive T cells were subjected to single-cell analysis using the 10x Genomics Chromium Single Cell 5′ Gene Expression and V(D)J library preparation workflow. Libraries were sequenced on an Illumina NovaSeq 6000 platform at the Advanced Technology Genomics Core, UT MD Anderson Cancer Center. Sequencing output was processed with the CellRanger software suite to perform sample demultiplexing, read alignment, barcode assignment, and reconstruction of CDR3 α and β chain clonotypes.

### Recombinant murinized TCR construction

To reduce the potential of mispairing with endogenous TCR chains, identified human TCR α/β sequences were murinized as previously described (30). Full-length, murinized TCRα and TCRβ genes were synthesized by GenScript USA, Inc. (Piscataway, NJ) and cloned into the pMSGV1 retroviral backbone following the strategy outlined by Hughes et al (31). TCR constructs incorporated a Furin cleavage site and a P2A self-cleaving peptide to link TCRα and TCRβ, facilitating efficient post-translational separation of the two chains (32).

### Generation of HER2 mutant-specific TCR-Ts

For retroviral production, 293GP packaging cells were seeded overnight on poly-Dlysine–coated 100-mm plates at 7×10^6^ cells per plate in complete DMEM. Cells were transfected with 6μg of the pMSGV1-TCR construct and 4.5μg of the RD114 envelope plasmid using Lipofectamine™ 3000 (Thermofisher, L00015). Viral supernatants were collected at 48and 72-hours post-transfection, pooled, and added to retronectin-coated (20 μg/mL, Takara Bio, T100B) non-tissue-culture-treated 24-well plates. Plates were centrifuged at 2,000g for 2 hours at 32 °C to facilitate viral loading. CD8⁺ T cells were isolated by negative selection using the Human CD8⁺ T Cell Isolation Kit (STEMCELL Technologies, 17953). Purified cells were activated with ImmunoCult™ Human CD3/CD28/CD2 T Cell Activator (STEMCELL Technologies, 10970) and 200 IU/mL recombinant human IL-2 and cultured for 48 hours in 50:50 medium (1/2 RPMI + 10% FBS, 1/2 AIM V). Activated T cells were transferred onto the virally loaded plates and spinoculated at 800g for 20 minutes at 32°C. This transduction step was repeated on two consecutive days to enhance efficiency. Transduced T cells were assessed 3–4 days later by flow cytometry for surface expression of the mouse TCR constant region and cognate tetramer binding.

### T cell immunoassays

<u>Functional avidity</u> was evaluated by measuring MIP-1β (CCL4) secretion following stimulation with mutant (MUT) or wild-type (WT) peptide. H1975-A*02:01 cells were pulsed with serial dilutions of peptide for 2 hours at 37°C, followed by washing to remove excess peptide. T cells were then added at a 1:1 effector-to-target (E:T) ratio and cocultured for 24 hours. Supernatants were collected, and MIP-1β concentrations were quantified using the Human CCL4/MIP-1β Quantikine ELISA Kit (R&D Systems, DMB00) according to the manufacturer’s instructions. Cytokine concentrations were determined by interpolation from a standard curve generated for each assay.

<u>Cytotoxicity</u> was assessed using the Incucyte live-cell imaging system (Sartorius). Target cells were seeded in flat-bottom 96-well plates at 5×10^5^ cells per well and incubated overnight. The following day, cells were pulsed with peptide (1μg/mL) for 2 hours at 37°C. Excess peptide was removed and replaced with fresh medium prior to addition of T cells at E:T ratios ranging from 20:1 to 1:1. Plates were placed in the Incucyte S3, and images were acquired every 3 hours for 36 hours. Cytotoxicity was quantified by longitudinal measurement of GFP^+^ target cells using Incucyte Basic Analyzer software (Sartorius). For tumor serial rechallenge experiments, T cells were harvested and transferred every 48 hours to freshly seeded and peptide-pulsed target cells. A total of three sequential rechallenges were performed.

For <u>co-receptor dependency</u> assays, CD4⁺ T cells were isolated from HLA-A*02:01 positive PBMCs using the EasySep™ Human CD4⁺ T Cell Isolation Kit (STEMCELL Technologies) according to the manufacturer’s instructions. Purified CD4⁺ T cells were transduced with HER2 mutant–specific T cell receptors (TCRs). Antigen-specific reactivity was evaluated by tetramer staining and by functional avidity assays as described above. IFN-γ production was quantified in culture supernatants using the Human IFN-γ ELISA MAX™ Deluxe Set (BioLegend, 430104), following the manufacturer’s protocol.

<u>Cross-reactivity</u> was evaluated by alanine scanning mutagenesis, in which each residue of the mutant peptide was individually substituted with alanine (glycine at positions were alanine was present). Peptides were pulsed onto target cells at a concentration of 1 μg/mL and co-cultured with T cells as described above. MIP-1β secretion was quantified in supernatants as previously described. Residues were considered essential if alanine substitution reduced T cell activation to <80% of the response induced by the mutant peptide. A binding motif was defined by assigning “X” to positions where alanine substitution did not reduce activation and the corresponding amino acid at positions where substitution resulted in reduced activation. Derived motifs were analyzed using ScanProsite to identify potential mimotopes, as previously described (33, 34).

For the <u>TIMING assay</u>, nanowell arrays on a chip were fabricated as previously reported (35). The chip was placed in a plasma chamber (Harrick Plasma Inc; Ithaca, NY) for two min and treated with PLL[20]-g[3.5]-PEG(2)/PEG(3.4)-biotin (50%) (SuSoS; Zurich, Switzerland). Target cells were incubated with peptide (10 µg/mL) for 1 hour. Target and effector cells were then washed three times with PBS before staining. We labeled the target and effector cells with PKH26 Red and PKH67 Green (Sigma-Aldrich), respectively. Cells were then washed three times in full media and resuspended at a density of 1 million cells/mL. Effector and target cells were loaded onto the nanowell arrays and resuspended in IMDM, 10% FBS, and Annexin V Alexa Fluor 647. The chip was imaged in BrightField, Alexa Fluor 488, TexasRed, and Cy5 using a Zeiss Axio Observer (Oberkochen, Germany) for 8 hrs over 80 time points. Image analysis and cell tracking were performed as previously reported (35).

For <u>Time-Lapse Flow Cytometry,</u> HER2-mutant specific TCR T cells were thawed and rested overnight in RPMI-1640 cell culture medium containing 10% (v/v) FBS, and 1% (v/v) penicillin/streptomycin (P/S) at 37 °C with 5% CO_2_. The following morning, cells were barcoded with laser particle optical barcodes as described previously (36) and stained with Live/Dead stain (Zombie R718, BioLegend), followed by a panel of releasable anti-CD8-VioGreen, anti-CD69-PE, anti-CD25-APC, and anti-PD-1-PEVio770 antibodies (Miltenyi Biotec), as well as anti-CD154Brilliant Violet 421 (Biolegend) and anti-CD137Brilliant Violet 605 (Biolegend). For phenotyping, cells were additionally stained with anti-CCR7Brilliant Violet 650, anti-CD45RA-FITC, antiCD45RO-APC, anti-LAG-3APC-Fire 750, and anti-TIM-3PE-Fire 700 antibodies (BioLegend). Samples were run on the cyclic flow cytometer built as previously described (36), collected and then resuspended in 1% (v/v) of REAlease reagent (Miltenyi Biotec) for 10 min at room temperature to remove the antibody panel. Next, the cells were stimulated with H1975-A*02:01 target cells preloaded with 1µg/ml either of cognate (HER2 Mutant) or noncognate (WT) peptide for the specified amount of time (0h, 14h, and 36h post stimulation) and re-stained with the same antibody panel. After 36h acquisition, cells were captured and restimulated second time either with H1975-A*02:01 target cells preloaded with 1 µg/ml either of cognate (HER2 Mutant) or non-cognate (WT) peptide, or PMA/I for 6h. Anti-CD107aPE-Dazzle 594 (BioLegend) antibody was added at the time of restimulation, and Brefeldin A and Monencin were added 1h post restimulation for the remaining 5h. Next, cells were stained with Live/Dead stain (Zombie R718, BioLegend), fixed and permeabilized using Cytofix/Cytoperm™ (BD) according to the manufacturer’s instructions, and stained for intracellular markers using anti-Granzyme B-PE-Cy7 (BioLegend), anti-IFNγ-VioGreen (Miltenyi Biotec), and anti-TNFαAPC antibodies (BioLegend). All samples were acquired by the cyclic flow cytometer at each time-point. Flow Cytometry Standard (FCS) 3.0 files were analyzed and gated using FlowJo software version v10.10.1. Two-dimensional embeddings were computed using Pairwise Controlled Manifold Approximation (37). Unsupervised clustering was performed using FlowSOM (38) implemented in the Python flowsom package. Sankey flow diagrams were visualized using SankeyMATIC (https://sankeymatic.com), where CD69+ cells were classified as “active” and CD69cells were classified as “inactive” at each measured time-point.

### Structural modeling

Three-dimensional structures of the TCR–pMHC complexes were modeled using TCRmodel2 with default parameters (39). Structural visualization and analysis were performed in PyMOL, including surface representation, prediction of polar contacts, and calculation of TCR chain centers of mass.

### Statistical Analysis

Statistical analyses were performed using GraphPad Prism (version 10.6.1; GraphPad Software). All experiments were performed in quadruplicate (n = 4 technical replicates per condition) unless otherwise indicated. Data are presented as mean ± SD (or SEM, if that is what you used). Paired or unpaired two-tailed Student’s t-tests were used as appropriate to compare two groups. P values < 0.05 were considered statistically significant. Exact P values are reported throughout the text where applicable.

## Data Availability

The data that support the findings of this study are available from the corresponding author upon reasonable request.

## Results

### Recurrent, clonal HER2 hotspot mutations define a targetable subset of NSCLC

To determine the prevalence of HER2 mutations across malignancies, we analyzed data from The Cancer Genome Atlas (TCGA) (40, 41). HER2 mutations were observed most frequently in bladder cancer (12%) and uterine endometrioid carcinoma (8%), while lower frequencies (2–6%) were detected in melanoma, esophagogastric cancer, cholangiocarcinoma, colorectal cancer, breast cancer, and non–small cell lung cancer (NSCLC) (**Supplementary Figure 1A**). The distribution of mutational hotspots varied by tumor type. Across cancers, S310F/Y represented the most common hotspot mutation, accounting for 13.1% of all HER2 mutations (**Supplementary Figure 1B**). In contrast, lung adenocarcinoma demonstrated a distinct hotspot profile, with kinase domain mutations such as G776delinsVC and A775insYVMA (or Y772dupYVMA) emerging as predominant alterations (**Supplementary Figure 1C**).

To further define hotspot prevalence and distribution in NSCLC, we examined data from the previously published Guardant360 cohort, comprising 1,719 patients with newly diagnosed HER2-mutant NSCLC in the United States (3). In this cohort, the four most frequent HER2 hotspot mutations were A775insYVMA (41.6%), S310F (10.7%), G776delinsVC (7.5%), and S310Y (4.7%) (**Figure 1A-B**). The consistent recurrence of these hotspot mutations across this cohort supports their feasibility as shared, mutationspecific therapeutic targets.

**Figure 1.**
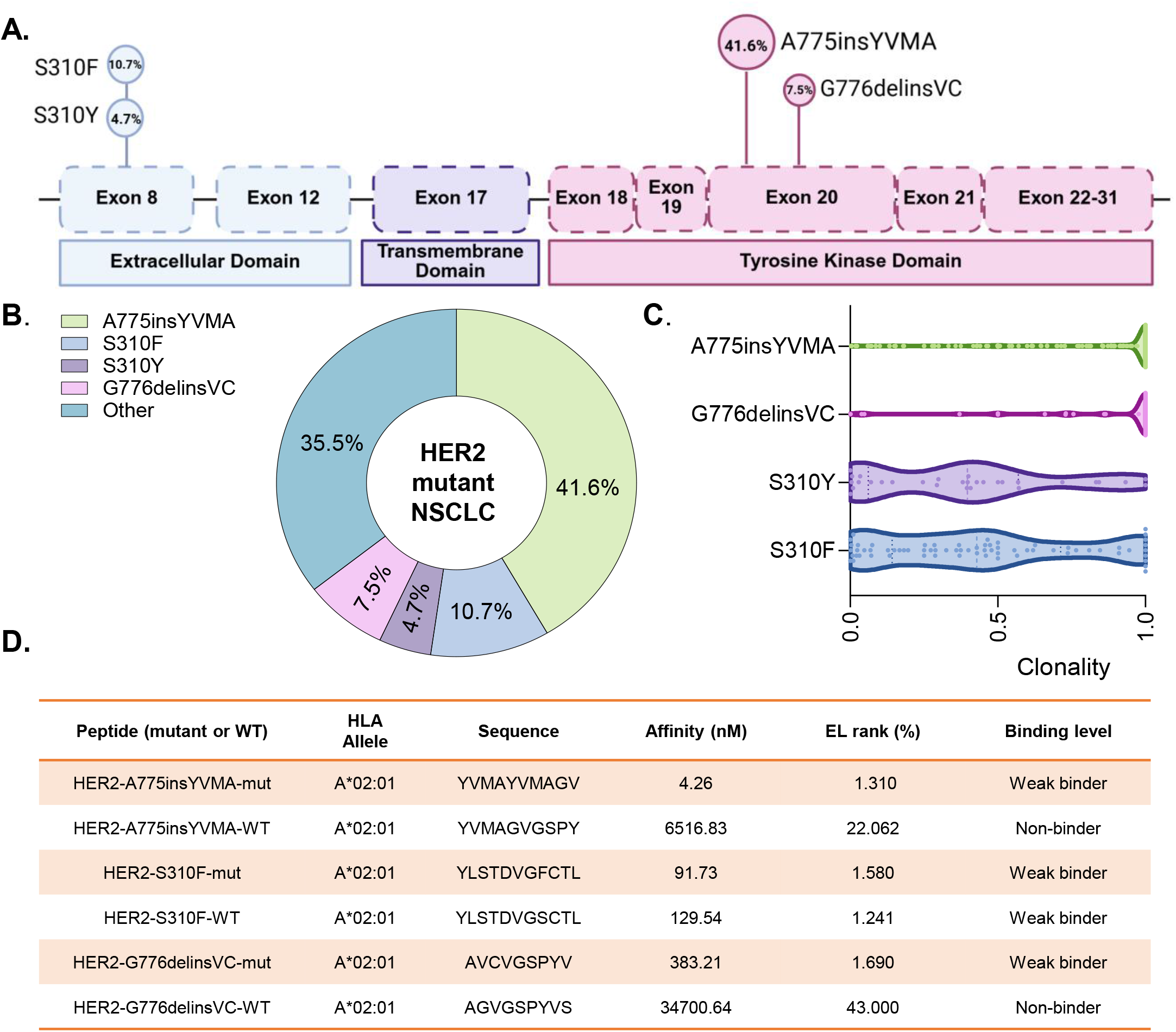
Distribution, clonality, and neoepitope characteristics of recurrent HER2 mutations in NSCLC. **A.** Location, frequency and **B.** distribution of recurrent HER2 mutations across HER2 protein domains. Percentages indicate the prevalence of each mutation among HER2-mutant NSCLC cases in the Guardant Health 360 (GH360) cohort**. C.** Clonality of the four most common HER2 mutations in the GH360 cohort. Each dot represents an individual sample. **D.** Predicted HLA-A*02:01 binding affinity of mutant and corresponding wild-type HER2-derived peptides. Data shown in panels A–C were derived from the previously published GH360 dataset (3).

Clonality was assessed using the ratio of variant allele frequency to maximum allele frequency. Kinase domain mutations demonstrated high clonality (ratio ∼ 1), suggesting their presence in most tumor cells while S310F/Y mutations exhibited intermediate clonality (ratio ∼ 0.5), (**Figure 1C**). Collectively, these data demonstrate that recurrent HER2 hotspot mutations in both the extracellular and kinase domains define an immunologically actionable subset of NSCLC.

We then selected predicted epitopes derived from the three most common HER2 mutations in NSCLC, which together account for approximately 60% of HER2 alterations. Candidate neoepitopes were prioritized based on predicted binding to HLA-A*02:01 using NetMHC 4.1, given the high prevalence of this allele in the U.S. population (42). Two of the selected mutations were indels, generating neoepitopes that differ from their corresponding wild-type sequences due to the introduced amino acid changes. Predicted HLA-A*02:01 binding was generally stronger for mutant peptides, whereas all wild-type peptides were predicted to be non-binders except for the S310F-derived epitope (**Figure 1D**).

### Identification and characterization of TCRs recognizing HLA-A*02:01–restricted HER2-Mutant neoepitopes

We previously established a workflow for the discovery and validation of T cell receptors (TCRs) from peripheral blood of healthy donors using *in vitro* stimulation (IVS) with predicted binding epitopes (**Figure 2A**) (43). Peripheral blood mononuclear cells (PBMCs) were differentiated into monocyte-derived dendritic cells, pulsed with candidate peptides, and co-cultured with autologous CD8+ T cells to generate antigen-specific populations. Following tetramer-guided sorting and a rapid expansion protocol (REP), enrichment of tetramer-positive T cells recognizing HER2-A775insYVMA, HER2-S310F, and HER2-G776delinsVC was observed, with marked increases in antigen-specific populations following REP (**Supplementary Figure 2A-C**). Single-cell RNA and TCR sequencing identified highly dominant clonotypes within each expanded antigen-specific population, with the most abundant clonotype comprising greater than 70% of cells in all three cultures (**Figure 2B-D, Supplementary Figure 2D-F**). Despite this clonal dominance, transcriptomic analysis revealed substantial phenotypic heterogeneity, with cells distributed across three major transcriptional states consisting of a proliferative Ki67^+^ effector population, a Ki67^−^ effector population, and a memory-like population (**Figure 2E-G**). Gene expression analysis revealed that these populations were characterized by expression of cytotoxic effector genes including GZMB, PRF1, GNLY, and NKG7, consistent with an antigen-experienced effector-like CD8^+^ T-cell phenotype (44), whereas inhibitory and exhaustion-associated markers were restricted to subsets of cells, supporting a predominantly effector-like phenotype with limited exhaustion features (**Figure 2H-J**). Finally, the CDR3α and CDR3β sequences of the dominant clonotypes were reconstructed, and the constant regions were murinized to reduce mispairing with endogenous TCR chains prior to downstream functional testing (30). TCR-T cells were generated from healthy donor PBMCs, with all constructs demonstrating robust expression following transduction (70–90% transduction efficiency).

**Figure 2.**
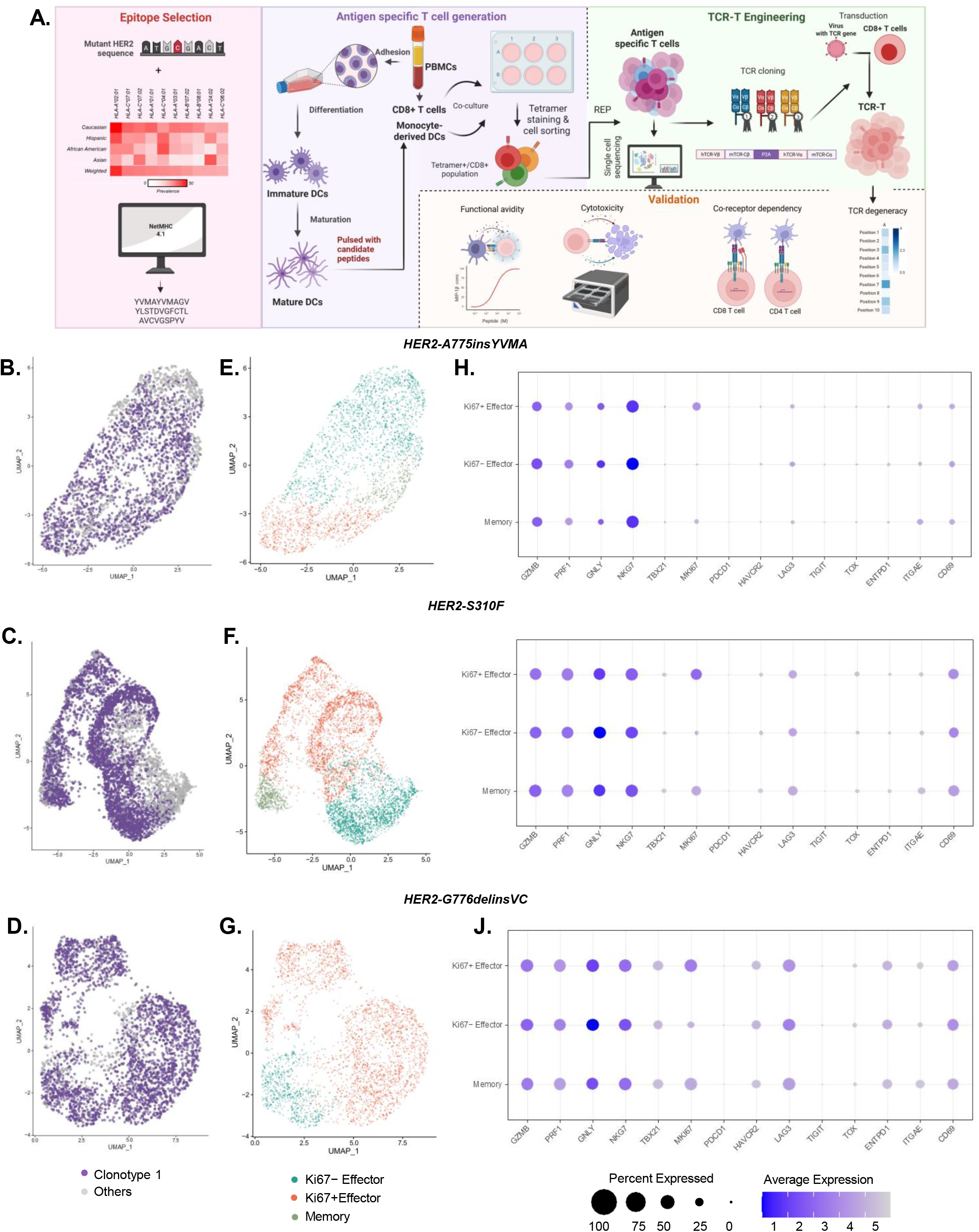
Identification and characterization of HER2 mutation-specific TCR clonotypes from healthy donor PBMCs. **A.** Schematic overview of the workflow used for HER2-specific TCR discovery and validation, including neoepitope selection, in vitro stimulation of healthy donor PBMCs, tetramer-guided enrichment of antigen-specific T cells, single-cell TCR sequencing, TCR engineering, and downstream functional characterization. Single-cell transcriptomic analysis of UMAP visualization of paired single-cell TCR sequencing data from tetramer-enriched T-cell populations specific for **B.** HER2A775insYVMA, **C.** HER2-S310F, and **D.** HER2-G776delinsVC with the most frequent clonotypes shown in purple and less frequent clonotypes shown in gray. Right UMAPs show cell-state annotation, demonstrating distribution of antigen-specific T cells across memory, proliferative (Ki67^+^ effector), and effector differentiation states. UMAP visualization of transcriptional states within **E.** HER2-A775insYVMA, **F.** HER2-S310F, and **G.** HER2-G776delinsVC antigen-specific T-cell populations. Dot plot representation of marker gene expression across transcriptional states for **H.** HER2-A775insYVMA, **I.** HER2-S310F, and **J.** HER2-G776delinsVC antigen-specific T cells. Dot size indicates the percentage of cells expressing each gene, and color intensity indicates average expression level.

### Functional Validation of HER2 Mutant–Specific TCR-T Cells

To assess antigen specificity, we first evaluated tetramer binding of TCR-engineered T cells. All TCR-T cells demonstrated selective binding to tetramers with their corresponding mutant epitopes, confirming mutation-specific recognition (**Figure 3A-C**).

**Figure 3.**
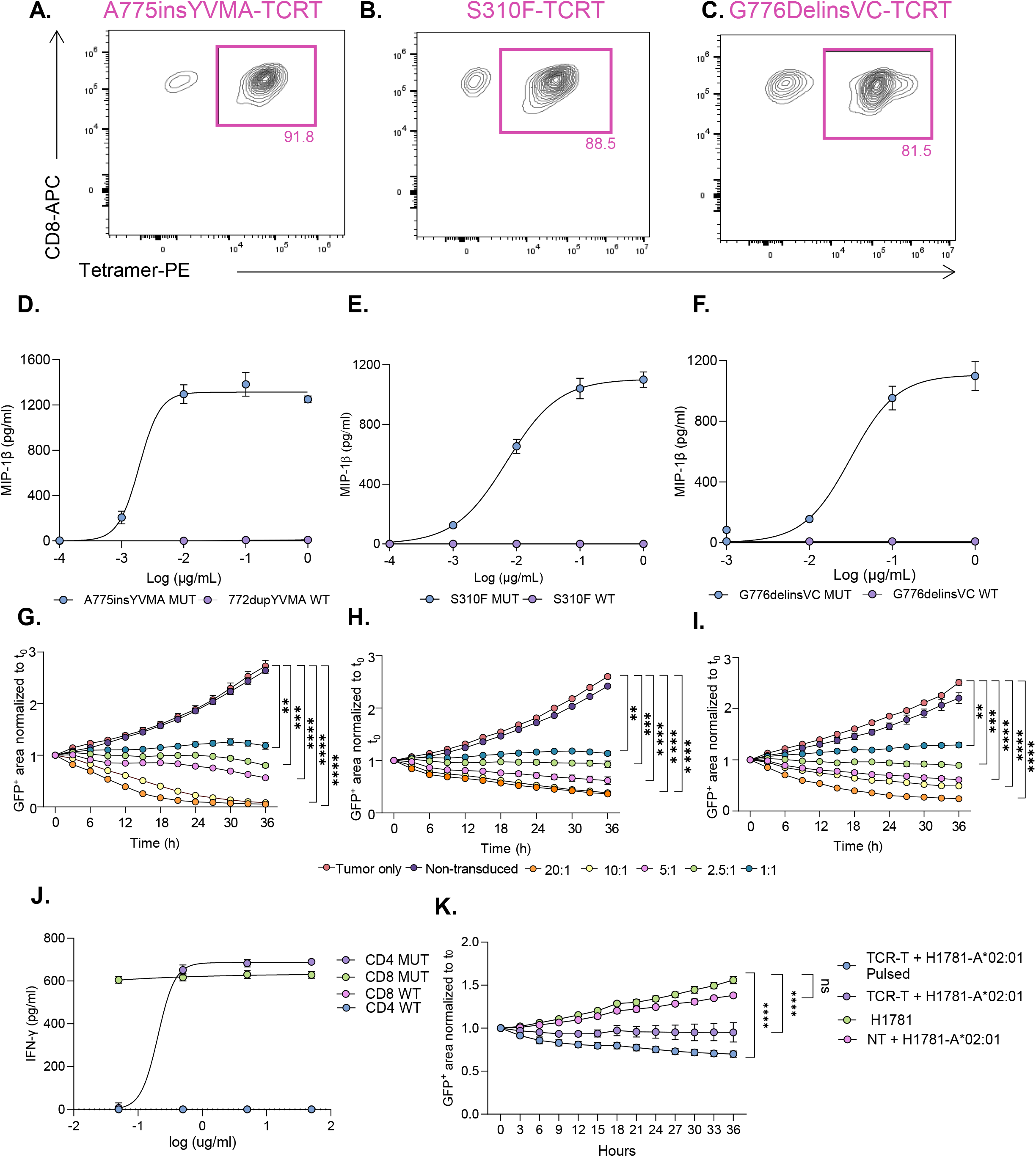
Generation and functional characterization of HER2 mutation–specific TCR-T cells. **A–C.** Representative tetramer staining of A775insYVMA, S310F, and G776delinsVC-specific TCR-transduced CD8+ T cells, demonstrating specific binding to cognate peptide–HLA complexes. Numbers indicate the percentage of tetramer-positive cells. **D–F.** Peptide titration assays of A775insYVMA, S310F, and G776delinsVCspecific TCR-T cells stimulated with target cells pulsed with mutant or corresponding wild-type HER2 peptides. Antigen-specific activation was measured by MIP-1β secretion. **G–I.** Live cell imaging cytotoxicity assays showing tumor cell killing by A775insYVMA, S310F, and G776delinsVC-specific TCR-T cells at the indicated effectorto-target (E: T) ratios. GFP area was normalized to baseline (t₀). **J.** IFN-γ secretion by G776delinsVC-specific CD4^+^ and CD8^+^TCR-T cells following stimulation with mutant or wild-type peptide. **K.** Cytotoxicity of endogenously processed HER2 G776delinsVC presented by H1781-A*02:01 cells. Tumor cell growth was monitored by normalized GFP area following coculture with G776delinsVC-specific TCR-T cells. Data are presented as mean ± SEM. Statistical significance is indicated as shown (*P < .05, **P < .01, ***P < .001, ****P < .0001).

T cell functional sensitivity was assessed via peptide dose-response by measuring T cell activation after serially diluting their cognate mutant or wildtype. All three TCRs exhibited robust dose-dependent responses to their cognate mutant peptides, while no reactivity was observed against the corresponding wild-type peptides (**Figure 3D–F**). T cell cytotoxicity was evaluated using live-cell imaging across multiple effector-to-target (E:T) ratios. All TCR-T cells mediated significant cytotoxicity at each ratio tested (P<0.05), demonstrating robust killing activity (**Figure 3G–I**).

Because CD8 co-receptor independence is indicative of high intrinsic TCR affinity, we next transduced each TCR into CD4⁺ T cells to assess functional activity in the absence of the CD8 co-receptor (33). The A775insYVMA and S310F-specific TCRs did not demonstrate detectable tetramer binding or functional activation in CD4⁺ T cells (**Supplementary Figure 3A-B**) In contrast, the G776delinsVC-specific TCR retained tetramer binding when expressed in CD4⁺ T cells and mediated dose-dependent IFN-γ production following stimulation with the HLA class I–restricted mutant epitope (**Figure 3J, Supplementary Figure 3C**). Moreover, the G776delinsVC-specific TCR mediated cytotoxicity against the lung cancer cell line H1781 (P<0.0001), which endogenously expresses the HER2 G776delinsVC mutation and was engineered to express HLAA*02:01 (**Figure 3K**), demonstrating recognition of endogenously processed and presented mutant antigen and supporting the translational potential of this receptor.

### Structural features of the S310F mutant epitope reveal cross-reactivity against closely related variant S310Y

*In silico* structural modeling of the S310F derived neoepitope revealed the aromatic chain at position 8 (phenylalanine) as a prominent exposed feature (**Figure 4A**). Since this residue corresponds to the oncogenic serine to phenylalanine substitution in HER2, we hypothesized that the closely related variant S310Y (serine to tyrosine) might also be recognized, as the tyrosine substitution preserves the aromatic exposed feature when modeled in the HLA-A*02:01 binding groove (**Figure 4B**).

**Figure 4.**
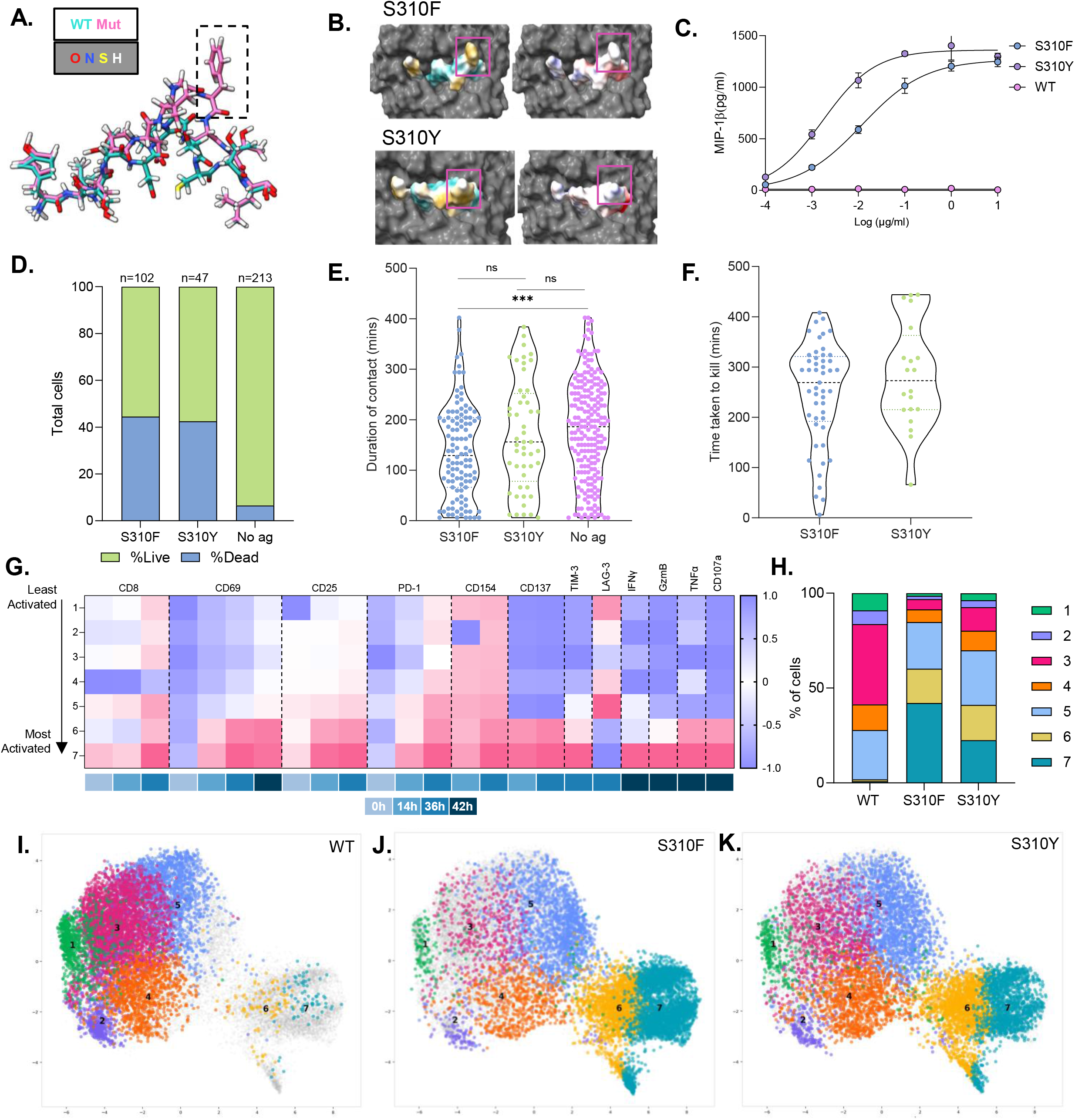
Structural and functional characterization of cross-recognition of HER2 S310F and S310Y neoantigens by a shared TCR. **A.** Structural overlay of the HER2 S310F mutant peptide (pink) and corresponding wild-type peptide (blue) demonstrating the prominent solvent-exposed aromatic residue introduced by the S310F mutation. **B.** Structural modeling of HER2 S310F and S310Y mutant peptides bound to HLAA*02:01. Insets highlight the mutant residue and local peptide–MHC conformation. Peptide surfaces are colored according to hydrophobicity (cyan, hydrophilic; gold, lipophilic) and electrostatic potential (red, negatively charged; blue, positively charged), with the HLA-A*02:01 molecule shown in gray. **C.** Peptide titration assay demonstrating activation of the S310-specific TCR in response to S310F, S310Y, and wild-type HER2 peptides. Antigen-specific activation was measured by MIP-1β secretion. **D.** Timelapse Imaging Microscopy In Nanowell Grids (TIMING) assay quantifying target cell survival following coculture with the S310-specific TCR-T cells. **E.** Duration of all T cell–target cell contacts measured by TIMING analysis. **F.** Duration of productive contacts preceding target cell killing. **G.** Heatmap showing median arcsinh-transformed expression of activation, inhibitory, and effector-function markers across FlowSOM meta-clusters (1–7) over time. Marker expression was normalized on a per-marker basis to visualize the dynamic range across clusters. Clusters 6 and 7 demonstrate progressive upregulation of activation markers, including CD69, CD25, and CD137, and are characterized by high expression of effector-function markers following 42-hour re-stimulation. **H.** Distribution of meta-clusters among HER2-specific TCR-T cells following coculture with untreated, wild-type HER2, S310F, or S310Y peptide–presenting target cells. Wild-type stimulated T cells were enriched for clusters 3 and 5, whereas S310F stimulation induced a marked expansion of activated/effector clusters 6 and 7. S310Y stimulation produced an intermediate phenotype, with increased representation of clusters 6 and 7 relative to wild-type conditions but to a lesser extent than S310F. **I-K**. Two-dimensional embedding of HER2-specific TCR-T cells following coculture with target cells presenting wild-type HER2, S310F, or S310Y peptides and subsequent FlowSOM clustering. Each point represents a single cell and is colored according to its assigned FlowSOM meta-cluster (clusters 1–7). The embedding identifies a major continuum comprising clusters 1–4, corresponding predominantly to non-activated and early-activated T-cell states, and a spatially distinct arm containing clusters 6 and 7, representing highly activated/effector T cells. Cluster 5 occupies an intermediate transitional state between these populations. **J.** Data are presented as mean ± SEM. Statistical significance is indicated as shown (ns, not significant;*P < .05, **P < .01, ***P < .001, ****P < .0001).

Functional assays confirmed a dose-dependent recognition of the S310Y derived neoepitope, along with cytotoxicity towards it (**Figure 4C & Supplementary Figure 4A**). Time-lapse imaging microscopy in nanowell grids (45) revealed distinct patterns of TCR-T engagement with each peptide, including differences in contact duration and time to target cell killing. However, both target cell populations were effectively eliminated by the TCR-T cells, highlighting the potency of these TCR-Ts against both variants (**Figure 4D-F & Supplementary Figure 4B**). Moreover, time lapse flow cytometry analysis of TCR-T cells after stimulation with the S310F and S310Y peptides revealed sustained upregulation of activation and effector markers (CD25, CD137, CD69, GzmB and CD107a) (**Figure 4G-J)**. Although both neoepitopes induced similar activation programs, differences in cluster representation were observed, with S310F stimulation favoring enrichment of highly activated effector clusters, whereas S310Y stimulation maintained a greater proportion of intermediate activation states (**Figure 4K**). These findings indicate that a single TCR can mediate cross-recognition of structurally related HER2 mutations, thereby expanding the targetable patient population and underscoring the potential of using *in silico* derived structural epitope features to expand the number of mutations a TCR can target.

### Alanine scanning mutagenesis reveals the key residues for TCR binding and potential cross-reactive mimotopes

To understand the molecular basis for the specificity of the A775insYVMA-reactive TCR, structural modeling of the TCR–peptide–HLA complex was performed (**Figure 5A**). The predicted interaction interface revealed hydrogen-bonding interactions between CDR3α and epitope residues Y1 and Y5, suggesting a critical role for these positions in TCR engagement. To experimentally define the residues required for antigen recognition, alanine-scanning mutagenesis was performed, with glycine substitutions used at positions where alanine was present in the native peptide sequence. (**Figure 5B**). While substitu-tions at several positions were tolerated, replacement of key residues resulted in a marked reduction in T-cell activation, defining a minimal recognition motif required for productive TCR engagement (Y-x-M-x-Y-V-M-x(3)). Consistent with the structural model, residues Y1 and Y5 were among the key determinants of TCR recognition. This motif was subsequently used to scan the human proteome for candidate peptides with similar sequence characteristics using the ProSite tool (33, 46). Only one candidate epitope was identified (**Supplementary Table 1**) and failed to elicit detectable cytokine production compared with the cognate mutant peptide, indicating an absence of measurable cross-reactivity despite partial sequence similarity (**Figure 5C**).

**Figure 5.**
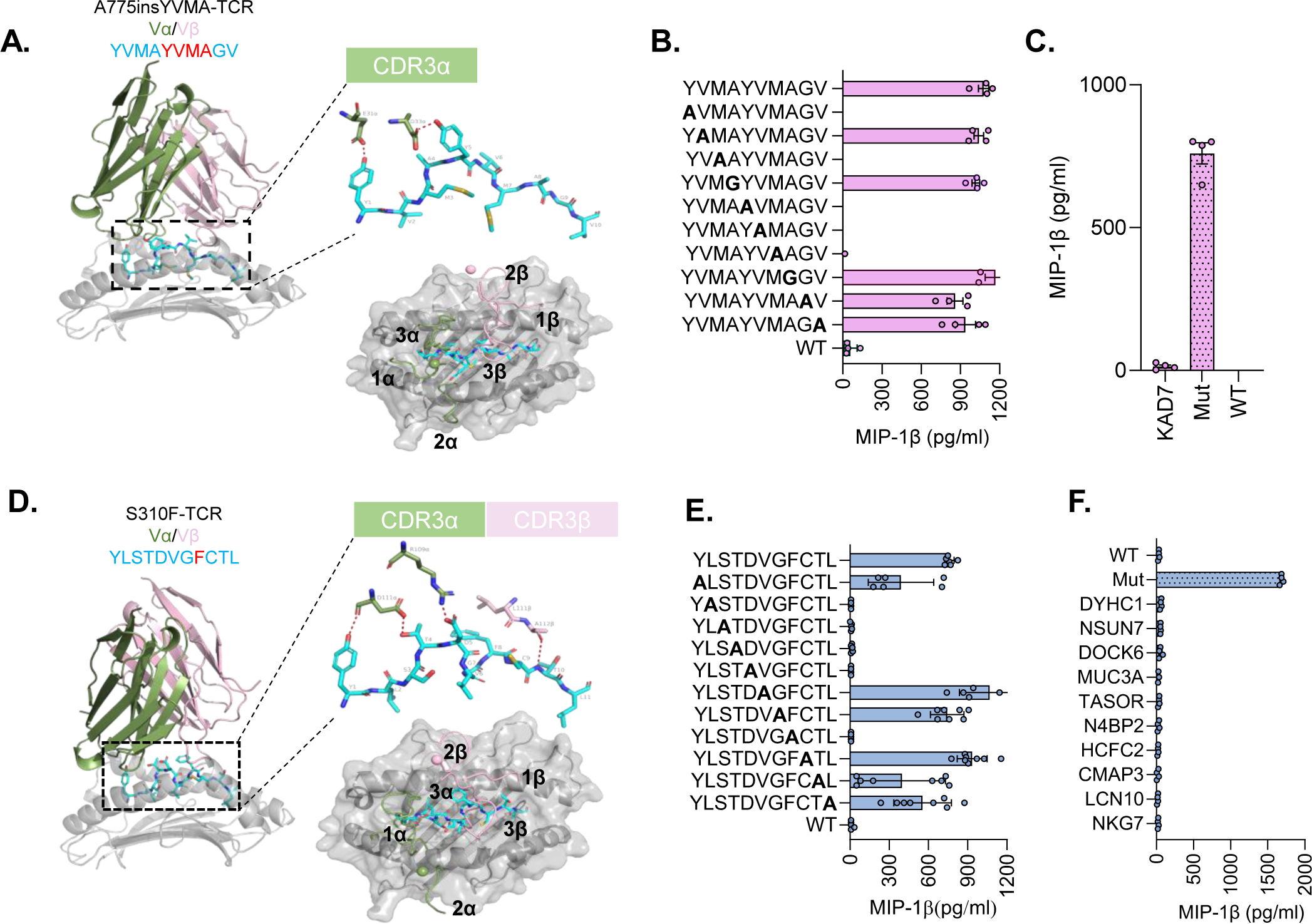
Mutation-specific TCRs exhibit high specificity with minimal off-target reactivity. **A.** Predicted structure of the A775insYVMA-specific TCR bound to the HER2 A775insYVMA peptide–HLA-A*02:01 complex. Insets highlight the TCR–peptide interface and demonstrate interactions with the CDR3α loop. **B.** Positional scanning of the HER2 A775insYVMA epitope (YVMAYVMAGV) to define residues required for TCR recognition. Individual peptide positions were substituted with alanine, or glycine when alanine was present in the native sequence, and T-cell activation was measured by MIP-1β secretion. **C.** Functional validation of candidate mimotopes identified using the minimal recognition motif derived from positional scanning. TCR activation was assessed following stimulation with peptide-pulsed target cells. **D.** Predicted structure of the S310F-specific TCR bound to the HER2 S310F peptide–HLA-A*02:01 complex. Insets highlight the TCR–peptide interface and demonstrate interactions involving both the CDR3α and CDR3β loops. **E.** Positional scanning of the HER2 S310F neoepitope (YLSTDVGFCTL) to define residues required for TCR recognition and generate a minimal recognition motif. T-cell activation was measured by MIP-1β secretion following stimulation with peptide-pulsed target cells. **F.** Functional evaluation of candidate selfpeptides identified using the minimal recognition motif derived from positional scanning. Minimal reactivity was observed against candidate self-peptides compared with the cognate mutant peptide, supporting a favorable specificity profile.

A similar approach was applied to the S310F-reactive TCR. Structural modeling of the TCR–peptide–HLA complex indicated interactions mediated through both the CDR3α and CDR3β loops, centered on the exposed phenylalanine residue generated by the oncogenic mutation (**Figure 5D**). Positional scanning identified a minimal recognition motif characterized by conserved residues L2, S3, T4, D5, and an aromatic residue at position F8 corresponding to the S310F substitution (**Figure 5E**). This motif (x-L-S-T-Dx(2)-[F/Y]-x(3)) was subsequently screened for candidate mimotopes (**Supplementary Table 2**); however, none of the 10 identified peptides elicited measurable T-cell activation compared with the cognate mutant peptide (**Figure 5F**), supporting a favorable specificity profile.

In contrast to the other neoantigen-reactive TCRs, positional alanine and glycine scanning of the G776delinsVC epitope did not reveal a clearly defined minimal recognition motif (**Supplementary Figure 5A**). Structural modeling of the G776delinsVC TCR– peptide–HLA complex demonstrated a focused interaction interface involving both TCR CDR3 loops and the mutant peptide (**Supplementary Figure 5B**). We therefore assessed reactivity against a panel of related HER2-derived peptides generated at the same mutational hotspot. While most variants failed to induce substantial IFN-γ production, the closely related G776delinsIC (p<0.0001) peptide elicited responses comparable to the cognate G776delinsVC neoepitope, and G776delinsIV induced an increase in T cell activation although this did not attain statistical significance (p=0.0649, **Supplementary Figure 5C**). These data suggest that this TCR can tolerate limited sequence variation among structurally similar insertion variants, while remaining highly selective against more divergent sequences and lacking wildtype reactivity. Collectively, these findings demonstrate that neoantigen recognition by these TCRs is governed by distinct structural interaction footprints that enable selective targeting of multiple clinically relevant oncogenic variants while maintaining a high degree of specificity against self-derived peptides, thereby broadening the potential patient population that could benefit from TCR-based therapies.

### Phenotype and persistence of HER2 mutant-specific TCR-T cells

Baseline phenotypic analysis demonstrated that all three TCR-engineered products were enriched for memory T-cell populations, with effector memory (Tem) cells representing the largest subset, alongside smaller fractions of naïve (Tn) and central memory (Tcm) cells (**Supplementary Figure 6A–B**). Within the CD45RA⁺CCR7⁻ compartment, a proportion of cells exhibited a CD45RO⁻ phenotype consistent with a terminally differentiated effector memory (TEMRA) phenotype across all three TCR products (**Supplementary Figure 6C**). Expression of the exhaustion markers TIM-3 and LAG-3 was low at baseline, with most cells lacking co-expression of these markers (**Supplementary Figure 6D–E**).

Furthermore, to evaluate the durability of T-cell function following repeated antigen exposure, TCR-engineered T cells were subjected to serial rounds of peptide stimulation and time-lapse flow cytometry (36).The A775insYVMA, S310F, and G776delinsVC-specific TCR-T cells each transitioned from predominantly inactive states at baseline into activated states following antigen stimulation and reacquired activated phenotypes upon antigen rechallenge (**Figure 6A–F**). Across all three TCR products, cells navigated through activated trajectories rather than accumulating in inactive populations. Upon restimulation with fresh peptide-expressing target cells, TCR-T cells maintained expression of cytotoxic markers, including granzyme B and CD107a, together with TNF-α and IFN-γ production, demonstrating preservation of polyfunctional effector states over time.

**Figure 6.**
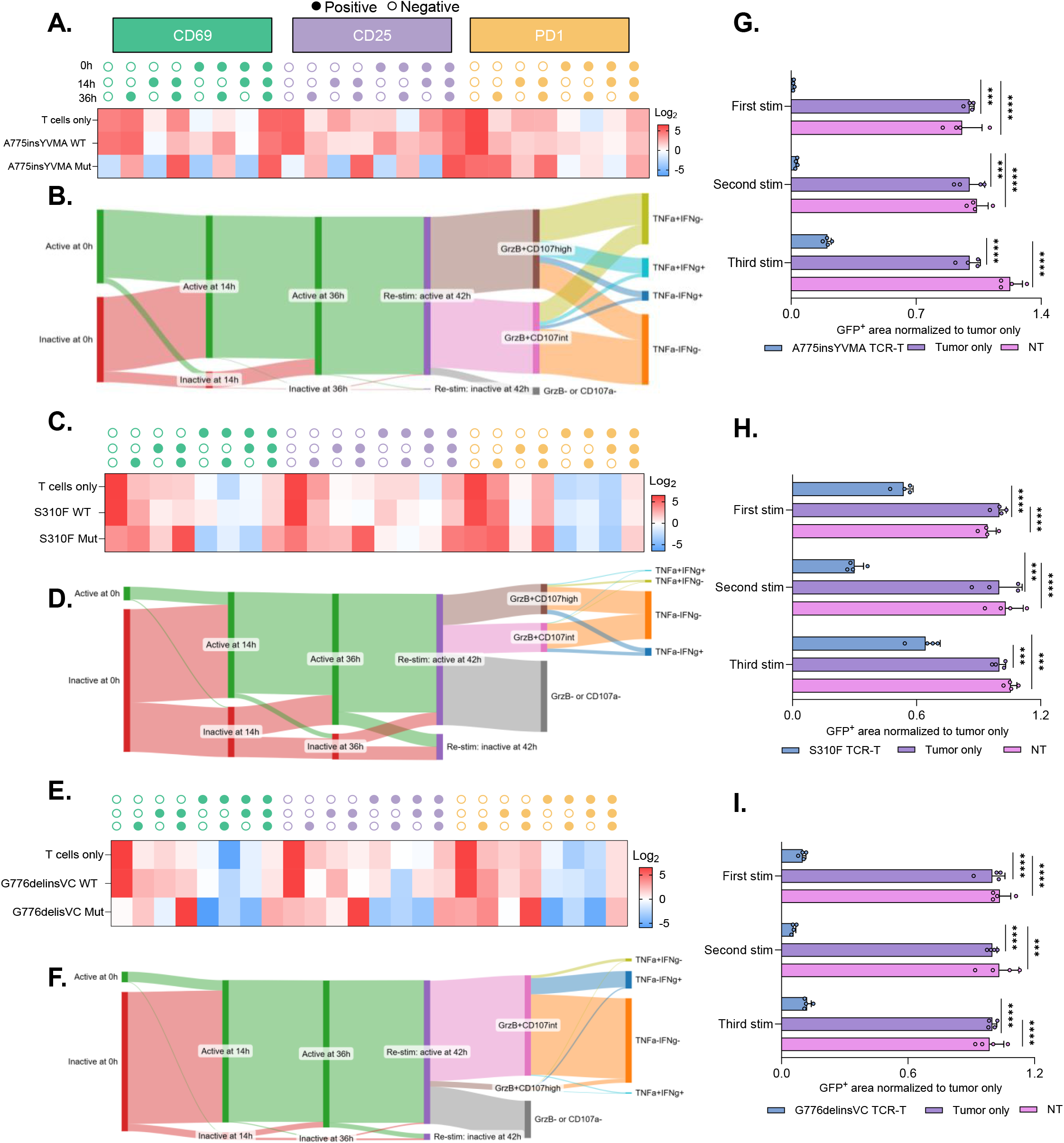
HER2 mutation-specific TCR-T cells maintain activation and antitumor function following repeated antigen exposure. **A–B.** Longitudinal activation profiles and state-transition analysis of A775insYVMA-specific TCR-T cells following serial rounds of stimulation with cognate mutant peptide, wild-type peptide, or no stimulation. Heatmaps depict the frequency of cells expressing activation markers (CD69, CD25, and PD-1) at 0,14 and 36 hours measured by time-lapse flow cytometry. State-transition analysis classified cells as active or inactive based on expression of activation markers (CD69, CD25, CD137, or CD154), followed by assessment of cytotoxic (Granzyme B and CD107a) and cytokine-producing (TNFα and IFNγ) phenotypes after restimulation. **C–D.** Longitudinal activation profiles and state-transition analysis of S310F-specific TCR-T cells. **E–F.** Longitudinal activation profiles and state-transition analysis of G776delinsVC-specific TCR-T cells. Serial tumor rechallenge assays demonstrating sustained antitumor activity of **G.** A775insYVMA**, H.** S310F, and **I.** G776delinsVC-specific TCR-T cells across three rounds of antigen encounter. Tumor burden was quantified by GFP⁺ area and normalized to tumor-only controls. Non-transduced (NT) T cells served as controls. Lower normalized GFP⁺ area indicates greater tumor killing. Data are presented as mean ± SEM. Statistical significance is indicated as shown (ns, not significant; *P < .05, **P < .01, ***P < .001, ****P < .0001).

To determine whether repeated antigen exposure affected tumor control, TCR-engineered T cells were subjected to serial tumor rechallenge cytotoxicity assays using live-cell imaging. All three TCR-T products retained the capacity to suppress tumor growth across multiple rounds of stimulation, although a gradual reduction in killing efficiency was observed with successive challenges (**Figure 6G, H, I**). Importantly, tumor control remained substantially greater than in tumor-only cultures throughout the experiment, indicating sustained functional competence despite repeated antigen exposure (p<0.001). Collectively, these findings demonstrate that HER2 neoantigen-specific TCRT cells retain activation potential, polyfunctional effector capacity, and tumoricidal activity following repeated antigen encounters, supporting their suitability for sustained antitumor responses.

## Discussion

HER2 mutations are established oncogenic drivers in 1–6% of NSCLC cases and are frequently associated with aggressive clinical behavior (1–4). Although HER2-targeted antibodies, kinase inhibitors, and antibody–drug conjugates have expanded treatment options, their benefit is limited by the inevitable acquired resistance (5–10). Consequently, there remains a critical need for therapeutic strategies capable of targeting HER2mutant tumors through mechanisms distinct from kinase inhibition. Recent advances in cancer immunotherapy have highlighted the therapeutic potential of targeting public neoantigens derived from recurrent oncogenic driver mutations, with TCRs directed against shared mutations in KRAS, PIK3CA, and TP53 demonstrating that a limited repertoire of mutation-specific receptors can potentially address large patient populations (22, 33, 47–50). Our findings extend this paradigm to HER2-mutant NSCLC and potentially to other cancers as well. Analysis of genomic datasets revealed that HER2mutant NSCLC is characterized by recurrent hotspot alterations, with A775insYVMA, S310F/Y, and G776delinsVC representing the most prevalent mutations. Importantly, these alterations frequently exhibited high clonality, particularly kinase-domain mutations, consistent with their role as early oncogenic driver events and suggesting a lower likelihood of immune escape through antigen loss. Together with the high prevalence of HLA-A*02:01 in North American populations (42), these features suggest that a relatively small panel of HER2-directed TCRs could provide therapeutic coverage for a substantial proportion of patients with HER2-mutant NSCLC. In this study, we developed and functionally validated three HLA-A*02:01–restricted TCRs directed against recurrent HER2 hotspot mutations (A775insYVMA, S310F and G776delinsVC) that collec-tively account for approximately 60% of HER2-mutant NSCLC, establishing HER2 hotspot mutations as actionable shared neoantigen targets.

Our findings extend prior observations demonstrating immune recognition of HER2derived neoantigens in NSCLC. Veatch and colleagues previously identified a naturally occurring CD4⁺ T-cell response directed against the HER2 YVMA exon 20 insertion (A775insYVMA) and isolated a class II–restricted TCR recognizing this mutation (26), providing important proof-of-principle evidence that recurrent HER2 driver mutations can serve as immunogenic targets in lung cancer. Building on this concept, we applied a previously developed platform to enrich antigen-specific T cells from healthy donor peripheral blood through *in vitro* stimulation with selected antigens, an approach that enabled the isolation of TCRs with high functionality and the capacity to mediate antigenspecific responses (43). Using *in silico*–predicted HER2 mutant neoepitopes with high predicted binding affinity for HLA-A*02:01, we identified and functionally validated HLAA*02:01–restricted class I TCRs targeting recurrent HER2 hotspot mutations. These TCRs demonstrated potent and highly specific recognition of their cognate mutant epitopes, with negligible detectable reactivity against corresponding wild-type peptides. Comprehensive cross-reactivity analyses incorporating alanine and glycine scanning mutagenesis, motif-guided mimotope identification, and functional testing against candidate off-target peptides further supported a favorable specificity profile. Collectively, these findings expand HER2-directed immunotherapy beyond previously described class II–restricted responses and establish a translational framework for the development of mutation-specific HER2 TCR-T cell therapies.

Several of the identified TCRs exhibited properties with important translational implications. The S310F-specific TCR demonstrated the ability to recognize the closely related S310Y variant, revealing a degree of cross-reactivity that broadens therapeutic applicability to this variant and other cancer types including bladder, cervical, colorectal, endometrial, and breast cancers (4). *In silico* structural modeling provided a mechanistic basis for this observation, showing that both mutated epitopes preserve a shared aromatic residue exposed at the TCR interface; notably, aromatic side chains have previously been associated with enhanced immunogenicity (51, 52). Consistent with these structural findings, the TCR mediated activation and cytotoxicity against both mutant variants, suggesting that some structurally related oncogenic mutations may be grouped into immunologically targetable families and potentially addressed by a single receptor without compromising specificity. In parallel, the G776delinsVC-specific TCR displayed several additional characteristics supporting its therapeutic potential, including retained functionality when expressed in CD4⁺ T cells, indicative of co-receptor independence and high intrinsic TCR affinity, and ability to recognize the G776delinsIC and G776delinsIV hotspots. Notably, CD4⁺ T cells have been shown to play important roles in mediating durable antitumor responses in adoptive T-cell therapies (23). Furthermore, this TCR was capable of recognizing and killing tumor cells endogenously expressing the HER2 G776delinsVC mutation demonstrating that HER2 mutation-derived neoepitopes can be naturally processed and presented. Collectively, these findings highlight opportunities to expand patient coverage through TCRs capable of recognizing structurally related mutant targets while supporting the feasibility of HER2-directed TCRT cell therapy.

Additionally, an important consideration for any adoptive T cell therapy is the durability of T-cell function following repeated antigen exposure. Using serial stimulation and rechallenge assays, we observed that all three HER2 mutant-specific TCR productsmaintained activation potential, cytokine production, cytotoxic activity, and tumor control across multiple rounds of antigen encounter. Although a gradual decline in killing efficiency was observed with successive challenges, T cells consistently retained substantial antitumor activity. These findings suggest that HER2 mutant-directed TCR-T cells possess the capacity to maintain functional competence following repeated antigen exposure, a characteristic that may be particularly important within the chronically stimulated tumor microenvironment.

Some limitations of this study warrant consideration. Although endogenous antigen recognition was confirmed for the G776delinsVC-specific TCR, validation of additional receptors was constrained by the limited availability of NSCLC models naturally harboring the relevant HER2 mutations in the context of appropriate HLA restriction. Future studies incorporating additional engineered systems or patient-derived models will be important to further evaluate endogenous processing and presentation of the targeted neoepitopes.

In conclusion, this study establishes recurrent HER2 hotspot mutations as actionable neoantigens in NSCLC and provides the first comprehensive characterization of class I HLA-restricted HER2 mutation-specific TCRs suitable for therapeutic development. By integrating genomic prevalence analyses, neoepitope prediction, structural modeling, and functional validation, we demonstrate that recurrent HER2 driver mutations can be selectively targeted by engineered T cells with high specificity and potent effector func-tion. As resistance to HER2-directed targeted therapies continues to emerge as a major clinical challenge, mutation-specific TCR-T therapies may offer a complementary and mechanistically distinct treatment strategy for patients with HER2-mutant NSCLC.

## CRediT Authorship Contribution Statement

**Amanda Montoya:** Conceptualization, Methodology, Investigation, Data curation, Formal analysis, Writing – original draft, Writing – review & editing.

**Hui Nie:** Conceptualization, Methodology, Investigation, Writing – review & editing.

**Peixin Jiang:** Formal analysis, Investigation, Data curation, Writing – review & editing.

**Jared Slone:** Methodology, Investigation, Data curation, Writing – review & editing.

**Yulia Shulga:** Conceptualization, Methodology, Formal analysis, Data curation, Writing – review & editing.

**Anika Patel:** Data curation, Writing – review & editing.

**Prashant Menon:** Methodology, Investigation, Data curation, Writing – review & editing.

**Isabella Polic:** Data curation, Methodology, Writing – review & editing.

**Emily Bontekoe:** Methodology, Writing – review & editing.

**Lingzhi Hong:** Methodology, Software, Data curation, Writing – review & editing.

**Minying Zhang:** Methodology, Writing – review & editing.

**Emane Rose Assita:** Investigation, Writing – review & editing.

**Sarah Forward:** Investigation, Writing – review & editing.

**Changsheng Xing:** Methodology, Writing – review & editing.

**Bo Jiang:** Methodology, Writing – review & editing.

**Drew C. Deniger:** Conceptualization, Methodology, Supervision, Writing – review & editing.

**Gregory A. Lizee:** Supervision, Writing – review & editing.

**Maura Gillison:** Supervison, Resources, Writing – review & editing.

**Navin Varadarajan:** Conceptualization, Methodology, Supervision, Resources, Writing – review & editing.

**Xiuning Le:** Conceptualization, Supervision, Writing – review & editing.

**Jianjun Zhang:** Conceptualization, Supervision, Writing – review & editing.

**Lydia Kavraki:** Conceptualization, Methodology, Supervision, Writing – review & editing.

**Sheldon J.J. Kwok:** Conceptualization, Methodology, Supervision, resources, Writing – review & editing.

**John V. Heymach:** Conceptualization, Methodology, Supervision, Resources, Writing – review & editing.

**Alexandre Reuben:** Conceptualization, Methodology, Project administration, Supervision, Funding acquisition, Resources, Writing – original draft, Writing – review & editing.

## Acknowledgements

The authors thank the MD Anderson Flow Cytometry Core Facility for technical assistance with flow cytometry studies. We also acknowledge the MD Anderson Single Cell Genomics Core for support with single-cell sequencing experiments, the MD Anderson Cytogenetics and Cell Authentication Core for cell line authentication services, and the Baylor College of Medicine MHC Tetramer Production Facility for the generation of MHC tetramers used in this work. Some figures were created with BioRender.

## Supplementary Figures and Tables

**Supplementary Figure 1. Landscape of ERBB2 (HER2) somatic mutations across TCGA cancers. A.** Frequency of ERBB2 alterations across tumor types in The Cancer Genome Atlas (TCGA). Bars represent the percentage of cases harboring ERBB2 alterations within each cancer type. **B.** Pan-cancer ERBB2 mutation spectrum. Somatic mutations were aggregated across all TCGA tumor types and mapped to ERBB2 amino acid positions. Mutation frequency is shown as a percentage of all ERBB2-mutant cases, with recurrent hotspot mutations labeled. **C.** ERBB2 mutation spectrum in TCGA lung adenocarcinoma (LUAD). Somatic mutations were mapped to ERBB2 amino acid positions and plotted according to their frequency within LUAD cases. Recurrent hotspot mutations occurring in ≥10% of ERBB2-mutant LUAD samples are labeled. Mutation frequencies and visualizations were generated from TCGA mutation annotation data using Python (pandas and matplotlib).

**Supplementary Figure 2. Enrichment and clonotypic composition of HER2 neoepitope-specific T-cell populations following rapid expansion.** Representative flow cytometric analysis of tetramer-positive CD8+ T cells specific for **A.** HER2A775insYVMA, **B.** HER2-S310F, and **C.** HER2-G776delinsVC following initial tetramer-guided sorting and after rapid expansion (REP). Red gates indicate tetramer-positive populations selected for downstream analysis. Clonotype frequencies identified by paired single-cell TCR sequencing of expanded **D.** HER2-A775insYVMA, **E.** HER2S310F, and **F.** HER2-G776delinsVC antigen-specific T-cell populations. Clonotypes are ranked according to relative frequency within each population.

**Supplementary Figure 3. Assessment of CD8 coreceptor dependency of ERBB2specific TCRs. A.** Representative flow plots showing tetramer (right) staining and mTCR (left) expression for the A775insYVMA-specific TCR in CD8⁺ and CD4⁺ T cells. Tetramer binding was observed primarily in CD8⁺ T cells, indicating CD8-dependent recognition. **B.** Representative flow plots showing tetramer (right) staining and mTCR (left) expression for the S310F-specific TCR. Tetramer-positive cells were predominantly CD8⁺, consistent with CD8-dependent recognition. **C.** Representative flow plots showing tetramer (right) staining and mTCR (left) expression for the G776delinsVC-specific TCR. Tetramer staining was observed in both CD8⁺ and CD4⁺ T cells, indicating CD8independent recognition. Blue gates indicate tetramer-positive cells and green gates indicate mTCR-positive cells.

**Supplementary Figure 4. Recognition and killing of ERBB2 S310Fand S310Ytarget cells by the S310F-specific TCR. A.** Representative wells from the TIMING assay showing interactions between S310F-specific TCR-transduced T cells and target cells pulsed with either the S310F or S310Y peptide over time. Images were acquired at the indicated time points**. B.** Live cell imaging cytotoxicity assay measuring killing of S310Y-pulsed target cells by S310F-specific TCR-transduced T cells at the indicated effector-to-target ratios. Green area was normalized to time 0 and monitored over 36 hours. Non-transduced (NT) T cells and target cells cultured without T cells served as controls.

**Supplementary Figure 5. Structural and functional characterization of the G776delinsVC-specific TCR. A.** Positional alanine and glycine scanning of the G776delinsVC neoepitope. Individual peptide positions were substituted with alanine or glycine and assessed for T-cell activation by measuring MIP-1β production. Responses are shown relative to the wild-type peptide and indicate the contribution of individual residues to TCR recognition. **B.** Structural model of the G776delinsVC TCR bound to the peptide–HLA complex. The model highlights interactions between the TCR CDR3α and CDR3β loops and the mutant peptide**. C.** Functional recognition of related HER2 G776 insertion variants by the G776delinsVC-specific TCR. TCR-transduced T cells were cocultured with target cells pulsed with the indicated peptides, and IFNγ production was measured by ELISA. The cognate G776delinsVC peptide and the closely related G776delinsIC variant elicited comparable responses, whereas G776delinsIV induced weaker activation and other variants failed to stimulate substantial cytokine production.

**Supplementary Figure 6. Baseline phenotypic characterization of ERBB2-specific TCR-engineered T-cell products. A.** Distribution of T-cell differentiation subsets within each TCR-engineered product. Cells were classified as naïve (Tn), stem cell memory (Tscm; CD45RA⁺CCR7⁺), effector memory (Tem), or central memory (Tcm) based on expression of CD45RA and CCR7. **B.** Memory phenotype distribution among CD45RA⁻ T cells. **C.** Expression of exhaustion-associated markers in TCR-engineered T-cell products. Frequencies of TIM-3⁺LAG-3⁺ cells and PD-1⁺ cells are shown for each TCR product, demonstrating low baseline expression of exhaustion markers prior to functional testing.

**Supplementary Table 1. Predicted cross-reactive human peptides for the HER2 A775insYVMA-specific TCR.** A ScanProsite motif search was performed using the minimal recognition motif identified for the HER2 A775insYVMA neoepitope. Candidate human peptides matching the motif were evaluated for predicted HLA binding using NetMHCpan. Listed peptides are derived from human proteins and include their corresponding UniProt identifiers, source proteins, peptide sequences, predicted HLA binding ranks (% Rank EL), and binding classifications.

**Supplementary Table 2. Predicted cross-reactive human peptides for the HER2 S310F/Y-specific TCR.** A ScanProsite motif search was performed using the recognition motif identified for the HER2 S310F/Y neoepitope. Candidate human peptides matching the motif were identified and evaluated for predicted HLA binding using NetMHCpan. Listed peptides are derived from human proteins and include their corresponding UniProt identifiers, source proteins, peptide sequences, predicted HLA binding ranks (% Rank EL), and binding classifications.

