## Supplementary figures and images for "HER2 mutation–derived neoantigens in NSCLC as actionable targets for TCR therapy"

### Supplementary Figure 1

*Supplementary Figure 1 . Montoya A. et al.*

**A.**

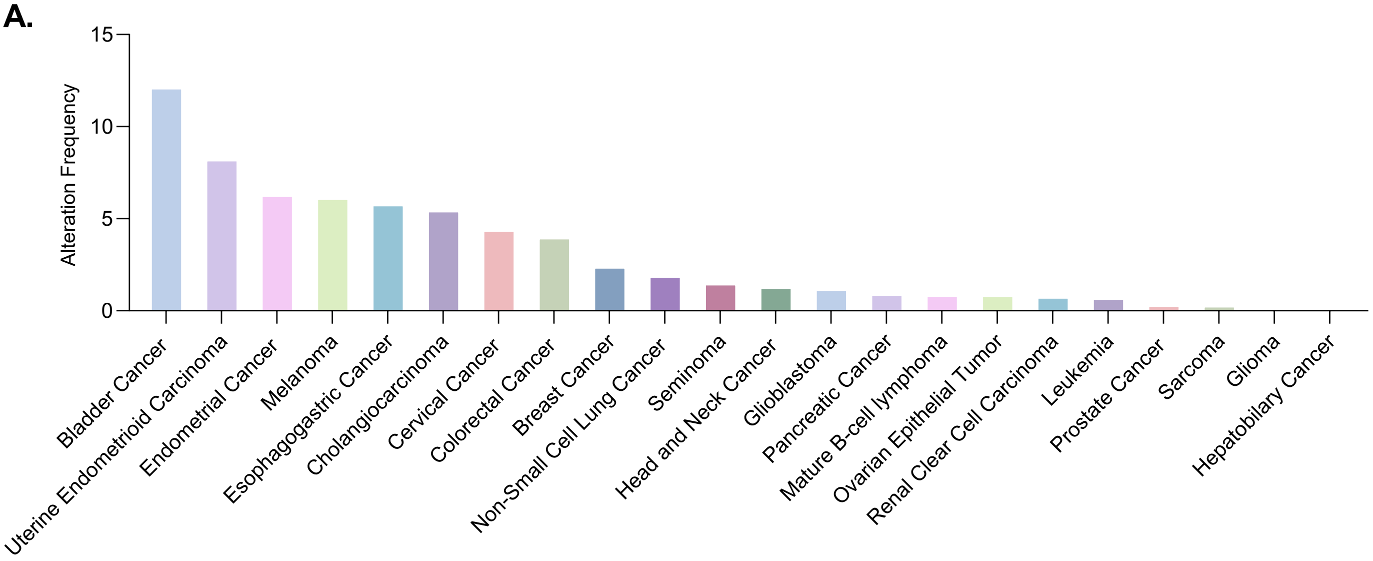


**B.**

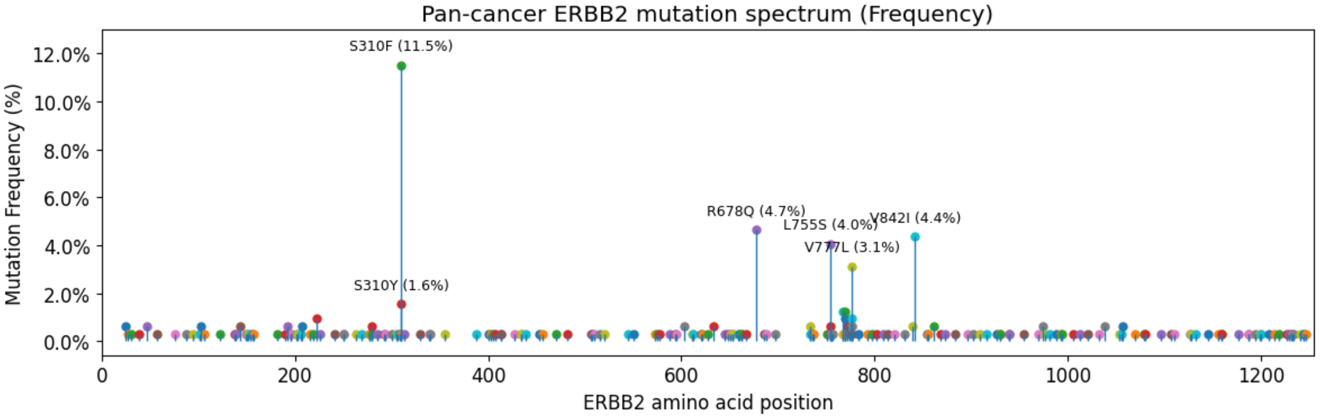


**C.**

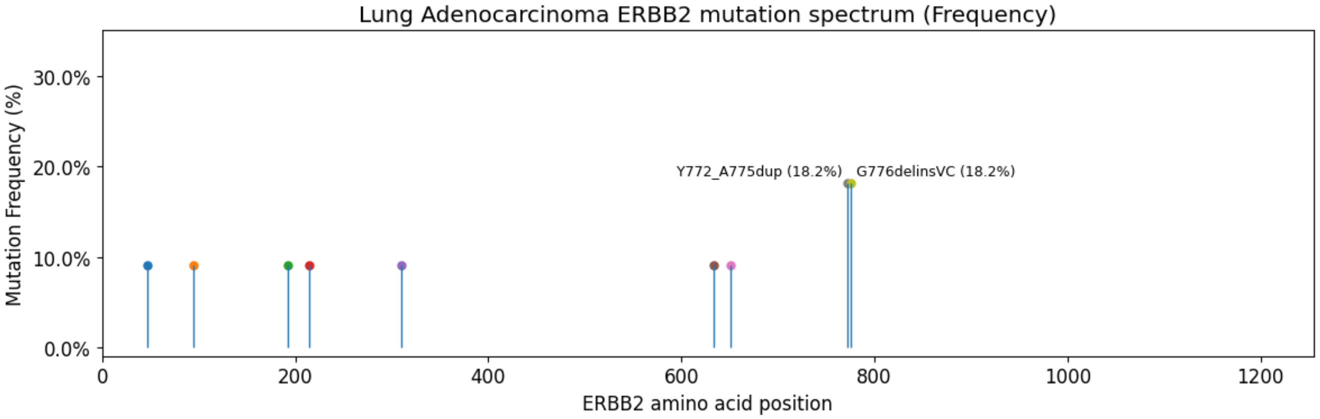

### Supplementary Figure 2

*Supplementary Figure 2. Montoya A. et al.*

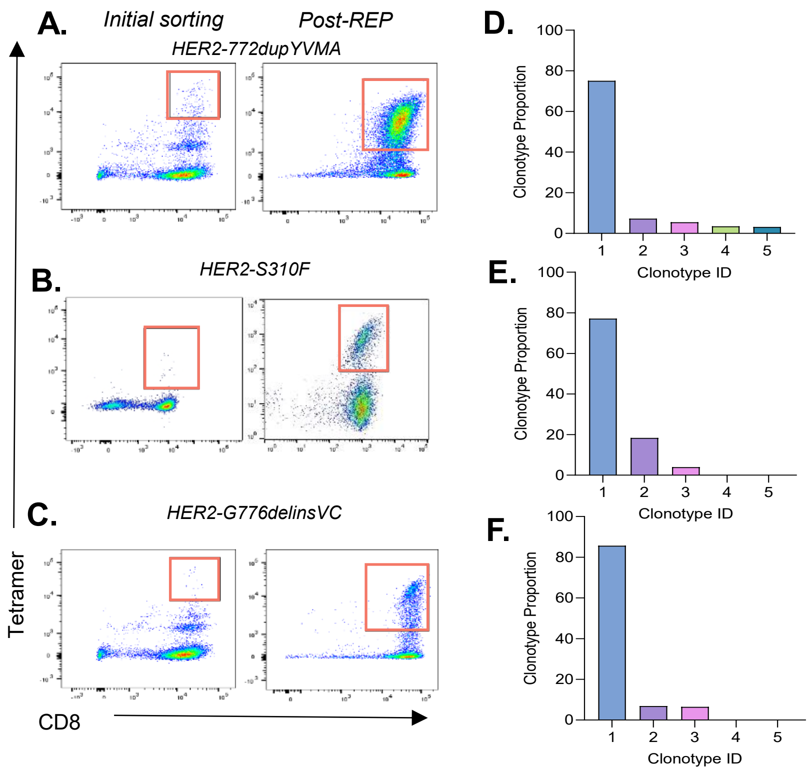

### Supplementary Figure 3

*Supplementary Figure 3. Montoya A. et al.*

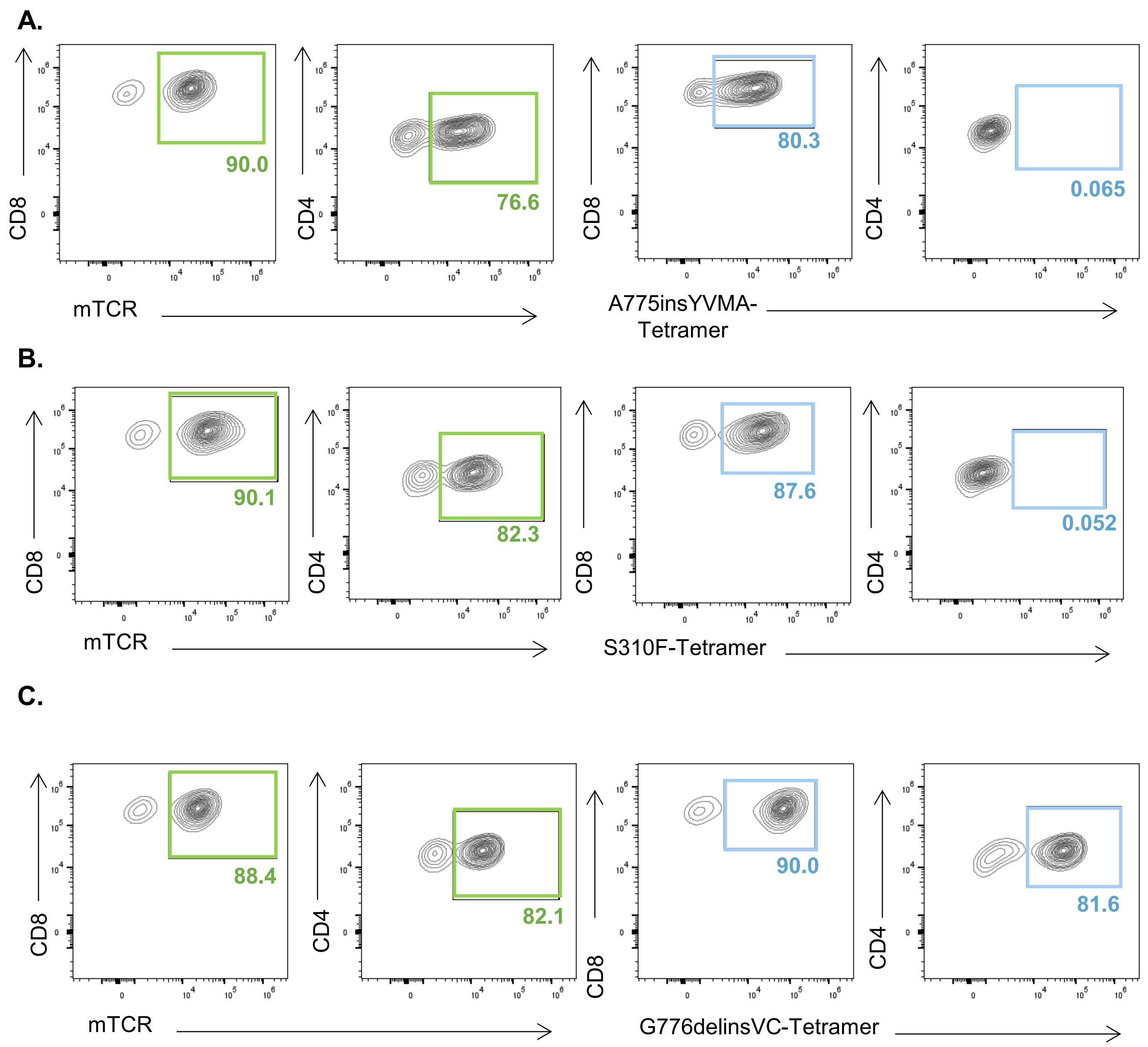

### Supplementary Figure 4

*Supplementary Figure 4. Montoya A. et al.*

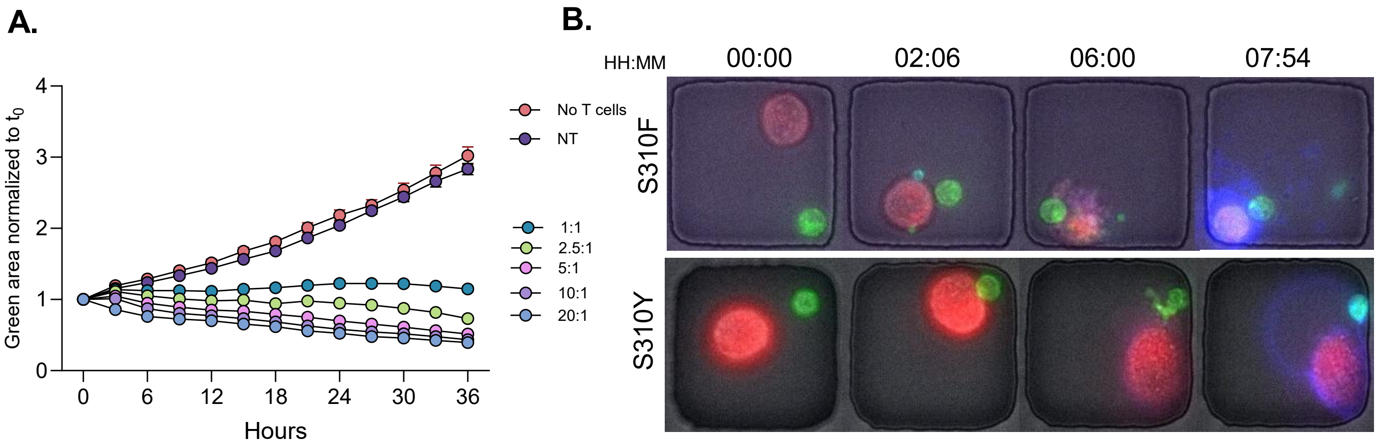

### Supplementary Figure 5

*Supplementary Figure 5. Montoya A. et al.*

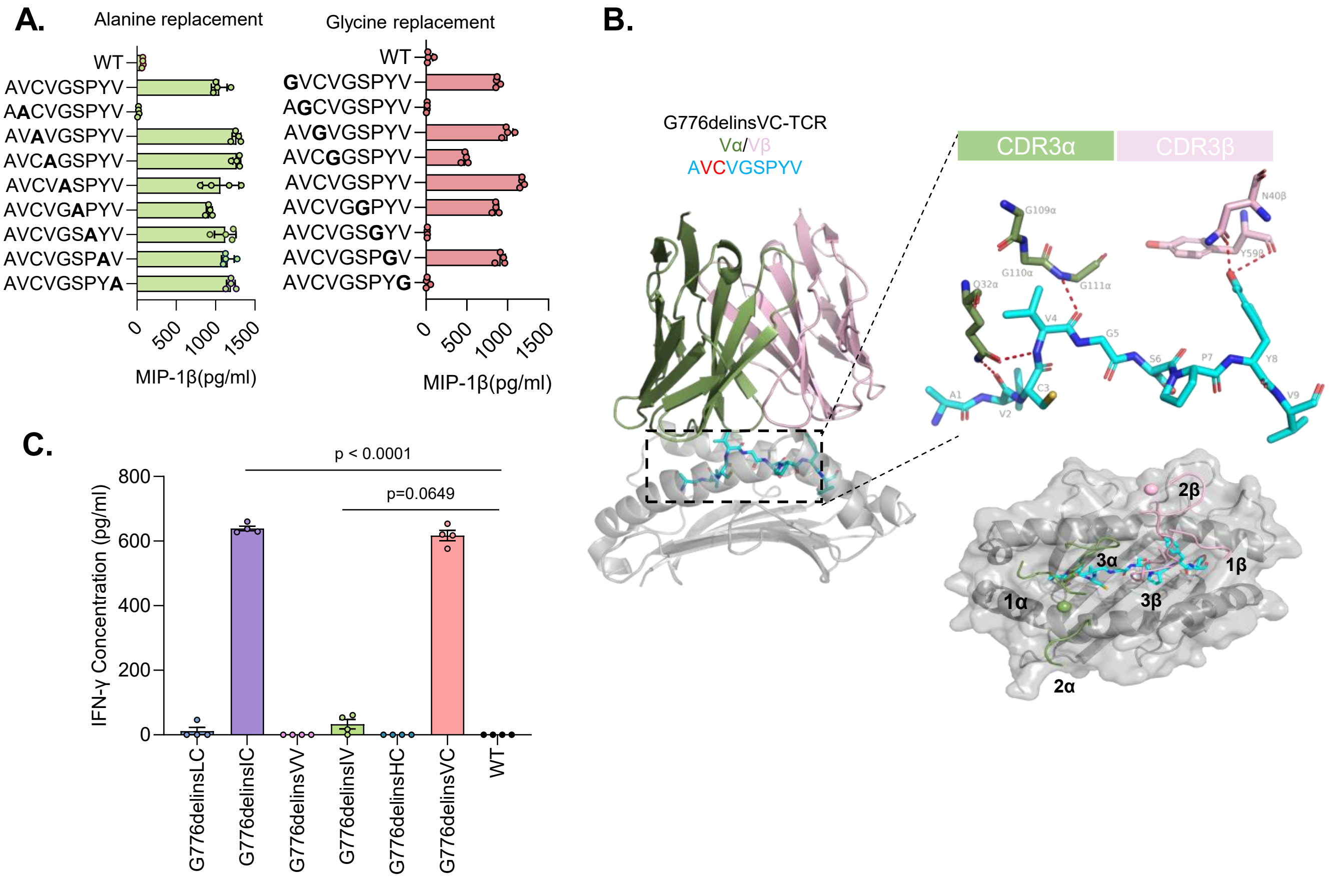

### Supplementary Figure 6

*Supplementary Figure 6. Montoya et al.*

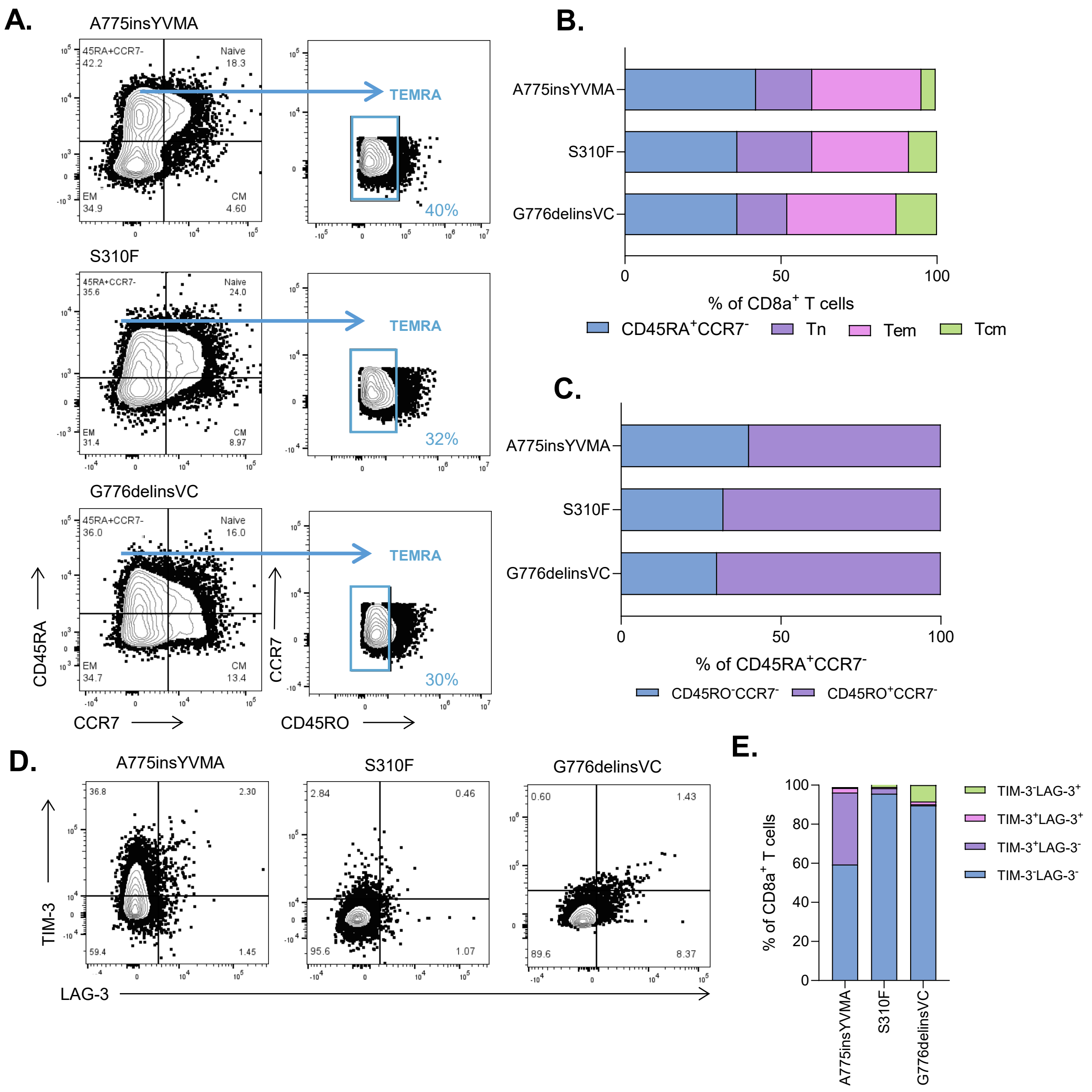
