## Supplementary Table 1 for "HER2 mutation–derived neoantigens in NSCLC as actionable targets for TCR therapy"

*Supplementary Table 1. Montoya A. et al.*

| *CROSS REACTIVITY ANALYSIS HER2 -A775insYVMA* | | | | | |
|---|---|---|---|---|---|
| **MOTIF:** | **Y-X-M-X-Y-V-M-X(3)** | | | | |
| **ScanProsite Output** | | | | | |
| **UNIPROT ID** | **IDENTIFIER** | **DESCRIPTION** | **EPITOPE** | **%Rank_EL** | **BIND LEVEL** |
| Q96M32 | **KAD7_HUMAN** | **Adenylate kinase 7** | **Y**L**M**T**YVM**PTL | 0.339 | SB |
