## Supplementary Table 2 for "HER2 mutation–derived neoantigens in NSCLC as actionable targets for TCR therapy"

*Supplementary Table 2. Montoya A. et al.*

| CROSS REACTIVITY ANALYSIS HER2-S310F/Y | | | | | |
|---|---|---|---|---|---|
| **MOTIF:** | **x-L-S-T-D-x(2)-[FY]-x(3)** | | | | |
| **ScanProsite Output** | | | | | |
| **UNIPROT ID** | **IDENTIFIER** | **DESCRIPTION** | **EPITOPE** | **%Rank_EL** | **BIND LEVEL** |
| Q8TCI5 | CMAP3_HUMAN | Ciliary microtubule-associated protein 3 | E**LSTD**KD**F**RKH | 45.333 | N/A |
| Q96HP0 | DOCK6_HUMAN | Dedicator of cytokinesis protein 6 | L**LSTD**HA**F**PYI | 2.838 | N/A |
| Q14204 | DYHC1_HUMAN | Cytoplasmic dynein 1 heavy chain 1 | V**LSTD**MI**F**NNF | 16.54 | N/A |
| Q9Y5Z7 | HCFC2_HUMAN | Host cell factor 2 | Q**LSTD**LP**Y**QAA | 3.706 | N/A |
| Q6JVE6 | LCN10_HUMAN | Epididymal-specific lipocalin-10 | V**LSTD**YS**Y**GLV | 4.666 | N/A |
| Q02505 | MUC3A_HUMAN | Mucin-3A | S**LSTD**IP**F**TTP | 6.387 | N/A |
| Q86UW6 | N4BP2_HUMAN | NEDD4-binding protein 2 | I**LSTD**DY**F**YIN | 10.736 | N/A |
| Q16617 | NKG7_HUMAN | Protein NKG7 | A**LSTD**FW**F**EAV | 1.536 | WB |
| Q8NE18 | NSUN7_HUMAN | Putative methyltransferase NSUN7 | Q**LSTD**KF**F**RME | 33.625 | N/A |
| Q9UK61 | TASOR_HUMAN | Protein TASOR | G**LSTD**DA**Y**EEL | 1.105 | WB |
